# Moonlight synchronous flights across three western palearctic swifts mirror size dependent prey preferences

**DOI:** 10.1101/2023.04.25.538243

**Authors:** Koen Hufkens, Christoph M. Meier, Ruben Evens, Josefa Arán Paredes, Hakan Karaardiç, Stef Vercauteren, Ann Van Gysel, James W. Fox, Carlos Miguel Pacheco, Luis P. da Silva, Sandra Fernandes, Pedro Henriques, Gonçalo Elias, Luís T. Costa, Martin Poot, Lyndon Kearsley

## Abstract

Recent studies have suggested the presence of moonlight mediated behaviour in avian aerial insectivores, such as swifts. At the same time swift species also show differences in prey (size) preferences. Here, we use the combined analysis of state-of-the-art activity logger data across three swift species, the Common, Pallid and Alpine swifts, to quantify flight height and activity responses to crepuscular and nocturnal light conditions. Our results show a significant response in flight heights to moonlight illuminance for Common and Pallid swifts, while a moonlight driven response is absent in Alpine swifts. Swift flight responses followed the size dependent altitude gradient of their insect prey. We show a weak relationship between night-time illuminance driven responses and twilight ascending behaviour, suggesting a decoupling of both crepuscular and night-time behaviour. We suggest that swifts optimise their flight behaviour to adapt to favourable night-time light conditions, driven by light responsive and size-dependent vertical insect stratification and weather conditions.

## Introduction

Relatively constant cycles of daylight and nocturnal darkness have shaped animal behaviour throughout evolutionary history (Aschoff 1989, Alerstam & Pettersson 1991, Guilford & Taylor 2014). Whereas most animals use photonic information from twilights as an external cue (i.e. zeitgeber) for daily activity, many crepuscular and nocturnal animals also use moonlight to trade-off behaviours associated with rest, reproduction, predation risk, and foraging (Smit *et al*. 2011, Appel *et al*. 2017, Evens *et al*. 2020, Hedenström *et al*. 2022, Pérez-Granados *et al*. 2022).

Nocturnal activity or changes in foraging tactics in response to moonlight through improved prey detectability have been typically attributed to nocturnal predators, such as nightjars, pelagic seabirds and owls (Cruz *et al*. 2013, San-Jose *et al*. 2019). In the case of insectivorous species, nightjars, for example, have been shown to increase flight activity and matching body temperature during bright moonlit nights, suggesting continued foraging throughout the night, when higher light levels facilitate prey detectability (Smit *et al*. 2011, Evens *et al*. 2020).

At the same time, many species of airborne insects show increased activity with available moonlight (Williams & Singh 1951, Bowden & Gibbs 1973, Danthanarayana 1986), determining the behaviour of predators. For example, Common Noctule bats (*Nyctalus noctula*) and Eleonora’s Falcons (*Falco eleonorae*) adjust their spatial activity during moonlit nights (Buij & Gschweng 2017, Roeleke *et al*. 2018). Not only does insect activity increase during moonlit nights, their distribution is also strongly vertically stratified, shaping food availability, with small insects flying higher than large ones (Helms *et al*. 2016). Aerial feeders heavily dependent on these insects select prey size according to their own body size (Collins *et al*. 2009). It follows that different species would find their prey at different altitudes.

Strong twilight ascents have been documented for the Common swift (*Apus apus*) using radar during the breeding season (Dokter *et al*. 2013, Nilsson *et al*. 2019). Similar observations have been made using multi-sensor geolocators across the non-breeding season for Alpine swifts (*Tachymarptis melba*, Meier *et al*. 2018). Both studies show strong increases in flight height and twilight activity. However, no strong connection was made between the duration of such twilight events and night-time (moonlight) illuminance. A study of Northern Black swifts (*Cypseloides niger borealis*) suggests that moonlit nights are used to continue foraging (Hedenström *et al*. 2022). During a lunar eclipse, briefly limiting moonlight, activity and flight altitudes of these black swifts decreased for the duration of the event only (Hedenström *et al*. 2022).

Recent research has highlighted changes in night-time flight activity and altitudes for Pallid swifts (*Apus pallidus*) (Hedenström *et al*. 2019, Kearsley *et al*. 2022). It was found that Pallid swifts locations coincide with dry locations on the northern (dry) side of the Inter-Tropical Convergence Zone (ITCZ), and support the notion that they might leverage foraging at a breaking nocturnal inversion during daytime. However, research on nightjars and black swifts put forward a new hypothesis of a predominantly moonlight driven response.

Despite indications of changes in night-time flight activity in both Common and Pallid swifts (Dokter *et al*. 2013, Hedenström *et al*. 2019, Kearsley *et al*. 2022), and documented night-time foraging during the breeding season in response to light pollution (Amichai & Kronfeld-Schor 2019), it remains uncertain if aerial insectivores such as the Common, Pallid or Alpine swifts extend their foraging behaviour into the night when natural illumination conditions are favourable.

Here, we use the combined analysis of state-of-the-art data logger data across three swift species to quantify flight height and activity responses to crepuscular and nocturnal light conditions and food availability. With small insects flying higher than large ones (Helms et al. 2016), it is reasonable to expect that *A. apus* and *A. pallidus*, which are smaller and differ in prey (size) preference (Collins *et al*. 2009), will likely fly at higher altitudes than *T. melba* during moonlit nights. We hypothesise that species with different feeding (prey) preferences show different responses to nocturnal light conditions.

## Methods

We deployed two types of data loggers at five sites across Portugal (2), Belgium (1), Switzerland (1) and Turkey (1), for Pallid, Common and Alpine swift respectively (Figure 1). The Portuguese Pallid swifts breed in the ceiling of a sea-side cave in the Serra da Arrábida Natural Park (38.47°N, 8.97°W, from hereon referred to as Arrabida), south of Lisbon and a second colony is housed in a municipal building in the town of Vila Nova de Famalicão (41.41°N, 8.52°W, from hereon referred to as Famalicão) approximately 20 km inland to the north of Porto. Belgian Common swifts nest in built-in nest boxes installed in a relatively new housing development (51.08°N, 3.73°E) along a dockside in the Ghent Voorhaven, Belgium (https://swifts.be/). The Alpine swifts nested under the roof of a historical town gate in Baden, central Switzerland, (47.47°N, 8.31°E), and in crevices of natural rocks along the coastline of the Pirasali island in southern Turkey (36.34°N, 30.53°E, from hereon referred to as Pirasali).

**Figure 1.**
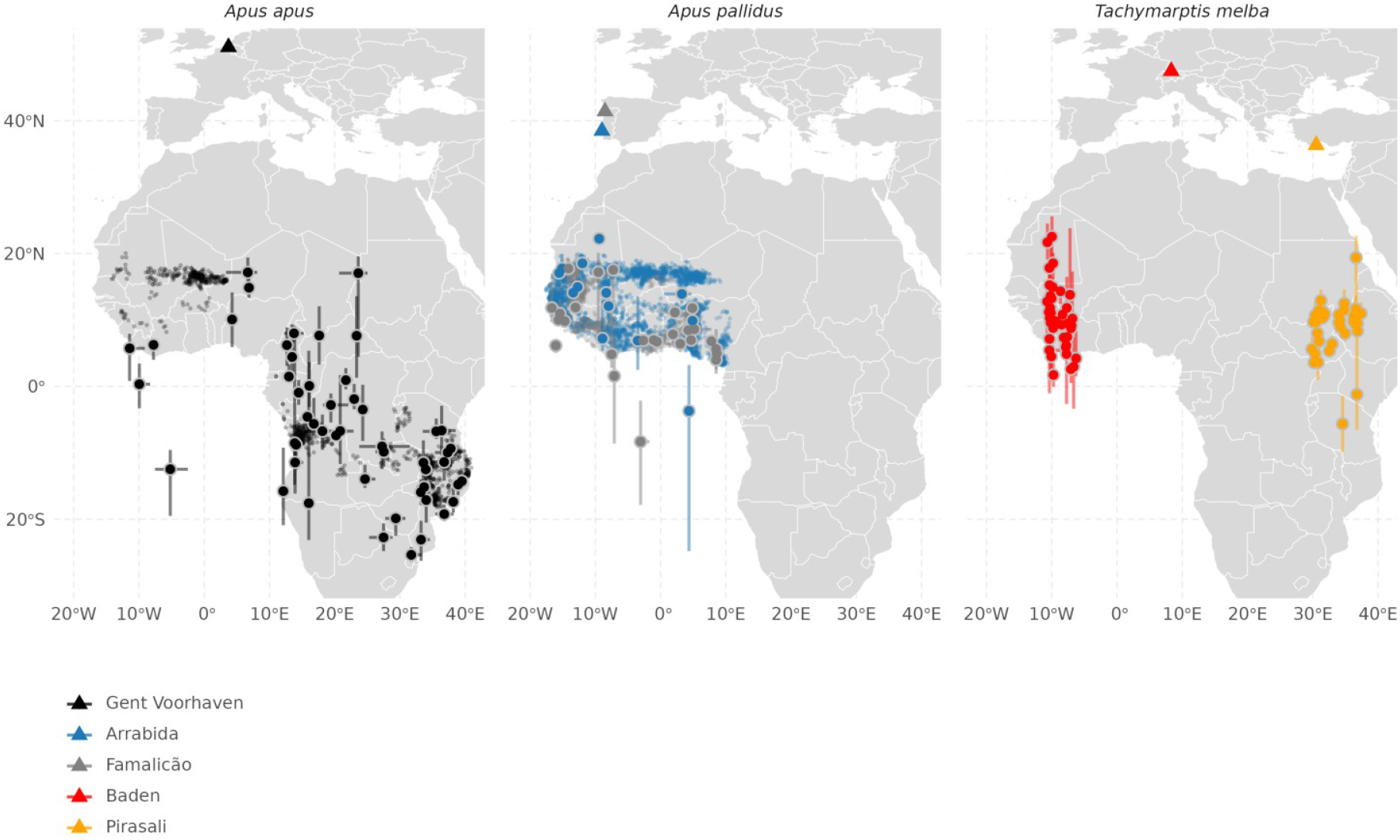
Overview map of both the GPS locations and the geolocator based estimated location data, and this for the five sites (colour triangles) and three swift species (panels: *Apus apus*, *Apus pallidus, Tachymarptis melba*). Small show GPS positions during stationary periods while positions during migration are omitted. Large show the median location of the cluster, with error bars showing one standard deviation around the mean. Colour dots correspond to site locations.

### Logger deployment

We used two lines of evidence based on separate animal tracking technologies across most sites. We used Multisensor Data Loggers (MDLs) for light level geolocation, pressure, and activity logging together with state of the art nanoFix micro-GPS loggers (Figure 2).

**Figure 2.**
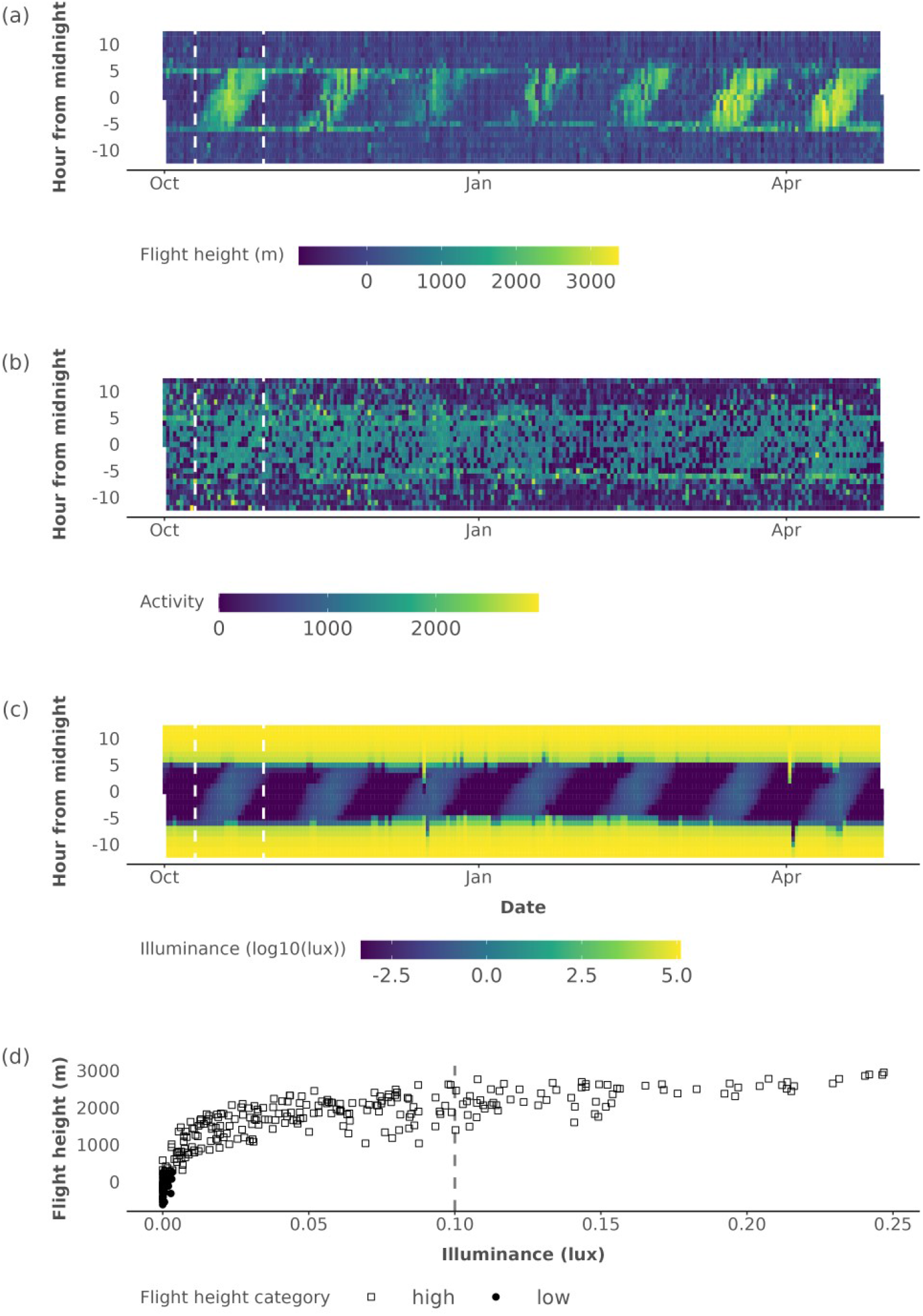
Continuous Multisensor Data Logger (normalised) flight height **(a)** and activity **(b)** measurements for Common swift (*Apus apus*) tag CC895 compared with modelled illuminance (log(lux)) values, **c**). In **(a-c)** time is centred on midnight, where negative hours indicate hours before midnight and positive values hours after midnight. And **(d)**, a scatterplot of the flight height changes in response to changing night-time illuminance values over the course of a lunar cycle of October 2021. Sampled dates of the scatterplot **(d)** are outlined by two white dashed lines in panels **(a-c)**, where flight height categories are marked with □ and, for high and low flights respectively. A grey vertical line specifies the moonlight illuminance value at which a ~50% probability of a high flight is exceeded (as modelled using a GLM).

Multisensor Data Loggers (MDL; Intigeo-BAR/CAR, Migrate Technology Ltd, Cambridge, UK) were deployed on 5 birds at the Arrabida site, and on 10 birds at the Famalicão site. Similarly, 9 data loggers were deployed on Voorhaven birds. We tagged eight Alpine swifts at the Swiss site and 26 at the Turkish Pirasali island with MDL from the Swiss Ornithological Institute. All MDL sensors logged light levels, pressure and temperature, while a number also logged activity (Appendix Table S1). Light intensity data was logged at five minute intervals while all other variables were recorded every 5 or 30 minutes (details see Appendix and Meier *et al*. 2020). Of a total of 58 MDLs were deployed across all sites, of which 29 were retrieved during the following breeding season (Appendix Table S3). Data was logged across four seasons from 2018 to 2021 (Appendix Table S1).

GPS tagging, with (≤ 1g) nanoFix micro-GPS loggers with solar trickle charging (PathTrack Ltd, Otley, UK) loggers, of 19 adult Pallid swifts across all Portuguese sites was carried out in parallel with the geolocator tagging. Swift diurnal positions and altitude were tracked throughout a full migration season, at regular six-hour intervals. Logging starts at 1h Greenwich Mean Time (GMT) for the Arrabida site, while Famalicão and Gent Voorhaven sites start logging at midnight (0h and/or 2h GMT). Data was logged over three seasons but with varying deployment dates (2019 - 2022, Appendix Table S2). Of 19 GPS loggers deployed across sites, 11 were retrieved during the following breeding seasons (Appendix Table S3). At the Famalicão site, two GPS loggers failed shortly after deployment and are therefore not included in our analysis. At the Gent Voorhaven site two GPS loggers were recovered but only a partial track was used for Tag21900 due to an age-related gradual battery failure (Appendix Table S1). Raw data was forwarded to PathTrack Ltd. for high precision processing.

Across all seasons we find overall recapture rates of ~55% and ~57% for MDL and GPS loggers, respectively, a value comparable with (long-term) ringed control groups (Costa & Elias 1998 and Appendix Table S3) although lower values were noted for the Alpine swifts (Meier *et al*. 2020). All loggers were deployed late in the breeding season to avoid breeding disturbance and followed the ongoing ringing protocol at the Portuguese colony (Costa & Elias 1998), and a similar approach at the Voorhaven site, i.e. using a full body harness with a 1mm wide flat braided soft terylene cord to avoid abrasion. Alpine swift protocols followed ringing and sensor deployment protocols as described in Meier *et al*. (2018, 2020), installing loggers at the end of the breeding season and collecting them before egg-laying started. Returning swifts were examined and no abrasion or skin damage was found.

### Data pre-processing

#### Location

GPS sensors provide near absolute positional accuracy in three dimensions (Kearsley *et al*. 2022). GPS data analysis separated stationary (area-restricted search) from migration movements using a Hidden Markov Model model. The non-breeding season analysed in this study was defined as all movements lower than 23.4° North, i.e. the Tropic of Cancer, constraining data to continental Africa below the Sahara desert. Swift movements were categorised as migration (corridor movements) and stationary (area-restricted search) using a two-state Hidden Markov Model (McClintock & Michelot 2018, HMM), with missing positions filled using continuous-time correlated random walk data (Johnson *et al*. 2008) to create a homogeneous six-hour time series (see Appendix, Kearsley et al. 2022). GPS positions were divided into “coastal” and “continental” positions, where a coastal position is defined as the two positions preceding and following a position above marine waters, with the coastline defined by 1:10m Natural Earth data (www.naturalearthdata.com). GPS time in GMT was corrected to local time using the longitudinal offset and centred on midnight.

Data were downloaded from MDLs and the approximate location of the Common and Pallid swifts was determined using geolocation by light (Lisovski & Hahn 2012). MDL data was constrained to the non-breeding season, determined by uninterrupted acceleration measurements and clear diurnal light patterns (Appendix Table S1). To mark twilight events we used a threshold of 1.5 on log-transformed and clock drift adjusted light level data. Clock drift was assumed to be linearly accumulating and was calculated from the difference between the on-board clock of the tag and a reference time. On a logger-by-logger basis we selected the most suitable sun-angle in half-degree steps by comparing derived positions with known GPS-based stationary locations (Appendix Table 1, Figure 1).

The median longitudinal difference between the breeding site and stationary positions were used to correct logger time to standardised local (solar) time, with a 4 minute per degree time difference. Location estimates were clustered in stationary and migration periods using a breakpoint analysis (i.e. changeLight in GeoLight, quantile parameter set to 0.9). For the processing of the Swiss and Turkish sites we refer to procedures described in Meier *et al*. (2020). We assigned the median longitude and latitude across a stationary cluster of dates to best approximate a swift’s location during that time. Missing location estimates (e.g. during equinoxes) were linearly interpolated from adjacent median cluster coordinates (Figure 1).

#### Flight altitude and activity

GPS logger processing can lead to spurious values due to limited reception which results in negative altitudes or locking values to 250m intervals if no clear altitude solution is found (personal communications PathTrack Ltd.). Therefore, we excluded all negative altitude values and those rounded to 250m increments. Flight altitudes were corrected for topography to flight height using NASADEM Digital Elevation Model (DEM) data (NASA 2020), with values below terrain level removed.

The MDLs accelerometer reads bursts along x and z-axis during a set measurement time interval (either 30 or 5 min, for Migrate Technology Intigeo-CAR or Swiss Ornithological Institute GDL sensors respectively, see Appendix). To ensure consistency across loggers we resampled the five minute GDL activity and pressure measurements to half-hourly intervals. In our analysis activity values should be seen as a proxy for motion. Atmospheric pressure recorded was converted to altitude (in m) using the international standard atmosphere (International Organization for Standardization ISO 2533:1975, see Appendix). We did not correct for local weather conditions as it only marginally affects altitude estimates (Hedenström *et al*. 2022).

Half-hourly MDL pressure and activity data was joined with the corresponding daily location estimates. GPS data highlighted considerable topographic differences across locations, e.g. Common swifts fly over a mean topography of > 500 m a.s.l., ~250 m higher than Pallid swifts (Appendix Table S3). Visually comparing absolute flight altitudes between species is therefore difficult and flight altitudes should be normalised for the underlying land surface elevation. Uncertainty on MDL positions limits correction of flight altitudes using DEM data. We accounted for day-to-day changes in underlying topography by subtracting the mean daytime altitudes on a day-by-day, using daytime (see below) flights as a fixed baseline (assuming relatively stable low flights during this period, e.g. Dokter *et al*. 2013). This method retains absolute diurnal altitude differences as a proxy for true flight heights, but might at times give rise to negative flight heights (Figure 2, Appendix Figures S1-S2).

We further divided flight heights into two classes using a threshold value based on the mean and standard deviation of daytime flight heights. night-time flight heights exceeding the mean daytime flight height plus one standard deviation, per individual (tag), were considered “high” flights and all others as “low” flights. This approach normalises responses between individuals (tags), allowing for a comparison across species irrespective of underlying topography and allows for an easy interpretation of statistical results.

#### Sun and Moon parameters

To facilitate further analysis, all data were divided into distinct periods according to the sun’s location. Here, “daytime” was defined as values with a sun angle above 0 degrees (above the horizon). “Dawn” and “dusk” were specified as times where the sun angle is between 0 degrees to 18 degrees below the horizon (the end of astronomical twilight) on the sun’s rising or setting direction. All other times are set to “night-time”.

Moon fraction and moon altitude are a poor substitute for moonlight exposure in animal studies (Kyba *et al*. 2017, 2020). We therefore improve upon general approaches using moon elevation and moon illuminated fraction by calculating moonlight exposure (illuminance) according to Janiczek & DeYoung (1987) for all positions using the ‘skylight’ R package (Hufkens 2022). We used MDL loggers’ median positions per stationary cluster (Figure 1), or absolute GPS position to calculate moonlight illuminance. Due to the large uncertainties on MDL light level geolocation, we did not account for local weather (cloud) conditions as this might weaken results rather than address any biases. We acknowledge that geolocation is not free of uncertainty. As such, estimated illuminance values are dependent on the accuracy of measured light derived geolocation, which might in term influence our modelling efforts. Yet, visual inspection of data centred on local midnight shows an expected centred twilight timing (Appendix S1-S4). We calculated an additional parameter for our crepuscular analysis, in particular the “twilight flight timing”. Twilight flight timing is defined as the timing of maximum flight height relative to timing of civil dusk or dawn, i.e. when the sun is 6° under the horizon (as calculated for a given location). Deviations of this twilight flight timing in response to moonlight illuminated night-time conditions would suggest an illuminance sensitivity in twilight behaviour.

### Statistical analysis

In our statistical analysis we test if flight activity and altitudes are influenced by moon illuminance. We divide our analysis into two parts. First we question if natural nocturnal light affects night-time flight height using both MDL and GPS sensors; and second, we test if nocturnal light affects twilight (dusk/dawn) activity, i.e. the effect on flight height when moving into or out of twilight periods.

To quantify the response of moon illuminance on the state of the flight height (high or low) we fitted a generalised linear mixed effects model (GLMM) with a binomial link function and using moon illuminance and lagged altitude, i.e. the altitude as measured at the previous time step, to account for temporal autocorrelation, and species as fixed effects (glmmTMB, Brooks *et al*. 2017). The tag is nested within species as a random intercept to account for species and bird specific behaviour. For all MDL measurements we only considered stationary periods. For the GPS data we used the same model. However, GPS measurements were measured at a low temporal frequency and therefore no correction for autocorrelation (i.e. lagged altitude) was applied. In our analysis we only retained continental positions. Coastal flight of predominantly Pallid swifts were excluded, as previously described coastal flight patterns (Kearsley *et al*. 2022) might skew the analysis.

The influence of night-time moonlight conditions on twilight behaviour, i.e. the flight behaviour moving into or out of twilight periods, is described by a linear mixed effects model with the twilight flight timing as response variable, moon illuminance is a fixed effects with tag nested within species as a random intercept. Similarly, we use a linear mixed effects model to quantify flight activity between species and the time of day (daytime vs. night-time). We modelled differences in flight activity as a function of time-of-day (day- vs. night-time) interacting with species as a fixed effect, while accounting for species and bird specific behaviour as random effects.

All statistics carried out in R (R Core Team 2023) using ‘glmmTMB’, ‘lme4’, ‘lmerTest’, and ‘marginaleffects’ packages, where visualisations were supported by the ‘tidyverse’, ‘patchwork’, ‘ggtext’ packages (Bates *et al*. 2015, Brooks *et al*. 2017, Kuznetsova *et al*. 2017, Wickham 2017, Arel-Bundock 2022, Pedersen 2022, Wilke 2022).

## Results

All swifts showed known migration patterns, with the Pallid swifts foraging in Western Africa during the non-breeding season (Hedenström *et al*. 2019, Norevik *et al*. 2019, Finlayson *et al*. 2021, Kearsley *et al*. 2022), while the Common swifts moved further to South-East Africa (Figure 1, Hedenström 2016). Alpine swifts migrated to either western or eastern Africa depending on their breeding sites, as previously described by Meier *et al*. (2020). We logged a total of 7235 GPS and 209 565 MDL positions with reliable data for statistical analysis. The MDLs geolocation patterns of stationary clusters correspond with those registered by GPS, despite the uncertainty on light level geolocation (Figure 1).

Distinct differences in maximum flight height are still noticeable between species (Figure 3a), despite normalising flight altitude data. Common swifts reach a maximum nocturnal flight height during a full moon of 1623 ± 873 m, while Pallid swifts on average flew at 929 ± 519 m and 896 ± 494 m for the Famalicão and Arrabida sites respectively. Alpine swifts did not show high nocturnal flights, with new moon night-time flights not differing much from full moon ones (331 ± 264 and 406 ± 334 m for Pirasali and 196 ± 183 and 253 ± 209m for Baden, Figure 3a).

**Figure 3.**
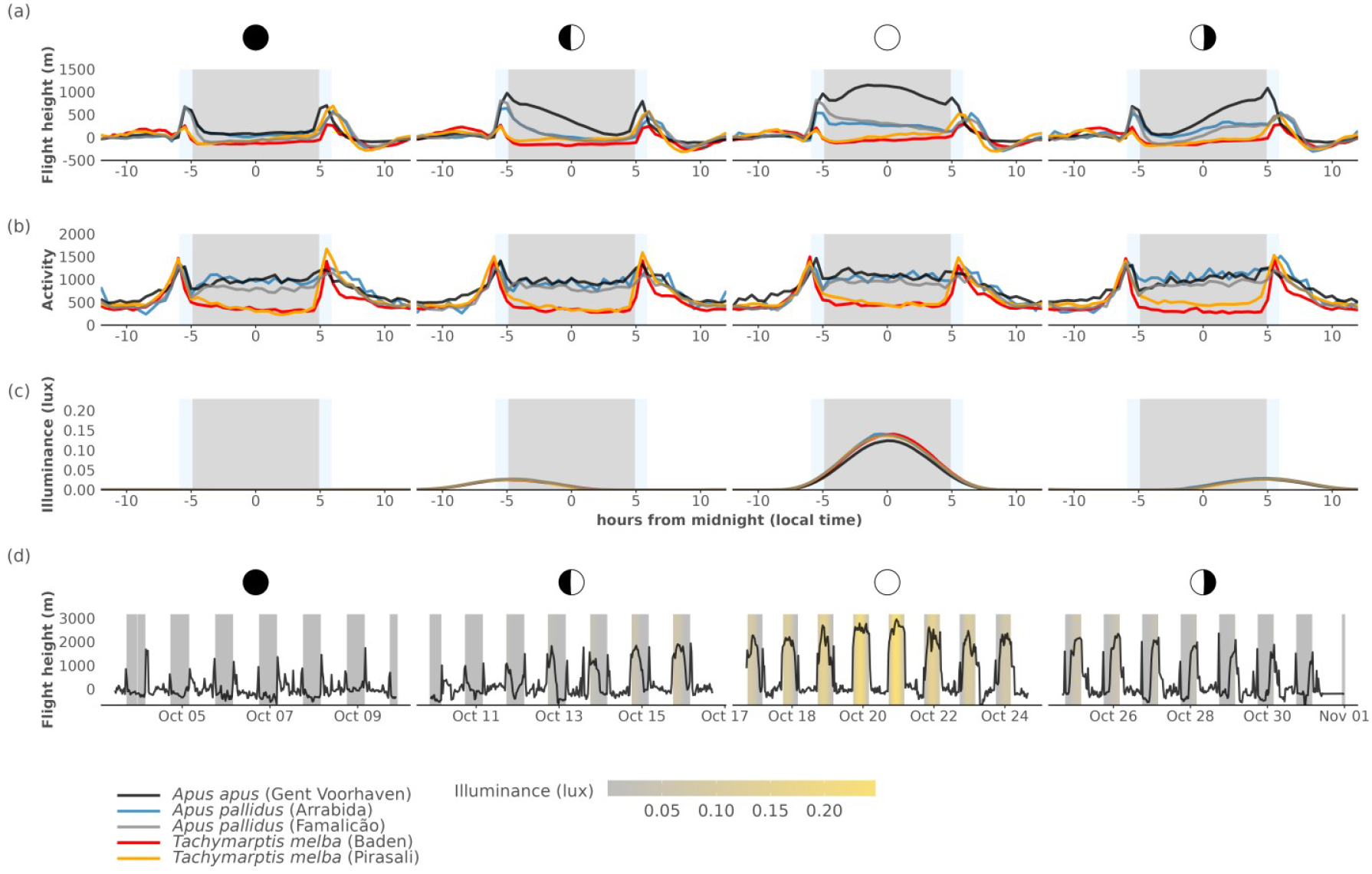
Average flight height (elevation in m, centred on a daytime mean of 0), the flight activity and derived moon illuminance (lux) centred on midnight during moon phases (a - c). Different sites and species are indicated by the colour of the full lines. The grey rectangle in plots **(a - c)** indicate the average astronomical night, where light blue rectangles approximate twilight as mean dusk and dawn ± 1 sd, respectively. In addition, a full time series of a lunar cycle **(d)** shows the progression of flight heights is shown in combination with moon illuminance values (daytime values are not shown, as orders of magnitude higher than the maximum illuminance of a full moon ~0.23 lux). We show time series for Common swift (*Apus apus*) tag CC895. Moon phases from left to right are indicated by their respective icons and are as follows: new moon, first quarter, full moon and last quarter. Activity **(b)** values of Alpine swifts (*Tachymarptis melba*) were scaled by a factor 12 to match the scale of other swifts (*Apus sp.*).

Across moon phases, we observe consistent patterns in changes in flight height and activity, and this for both GPS or MDL sensors. MDL flights are high when moonlight illuminance is high (Figure 3a, c and d, Appendix Table S3) for the Pallid and Common swifts. In contrast, Alpine swifts do not show a response to moonlight Figure 3a). Our MDL (GLMM) model showed a marginal R^2^ of 0.70 for the explanatory power of the fixed effects alone, with a significant effect of moon illuminance, temporal lag and species (Table 1). A similar model using GPS data, but omitting the lagged altitude response, shows that the explanatory power related to the fixed effects alone (marginal R^2^) is weak at 0.15, with a positive and significant effect of moon illuminance; yet no significant effect of species was found (Table 2). Our model results show different species responses, with Common and Pallid swift showing a strong response to moonlight illuminance (Figure 3), while Alpine swifts lack such a response (Figure 3a, Figure 4a). Two sensor technologies provide two lines of evidence and robust and consistent hypothesis support, where the high frequency of MDL data complements high accuracy of GPS based data.

**Figure 4.**
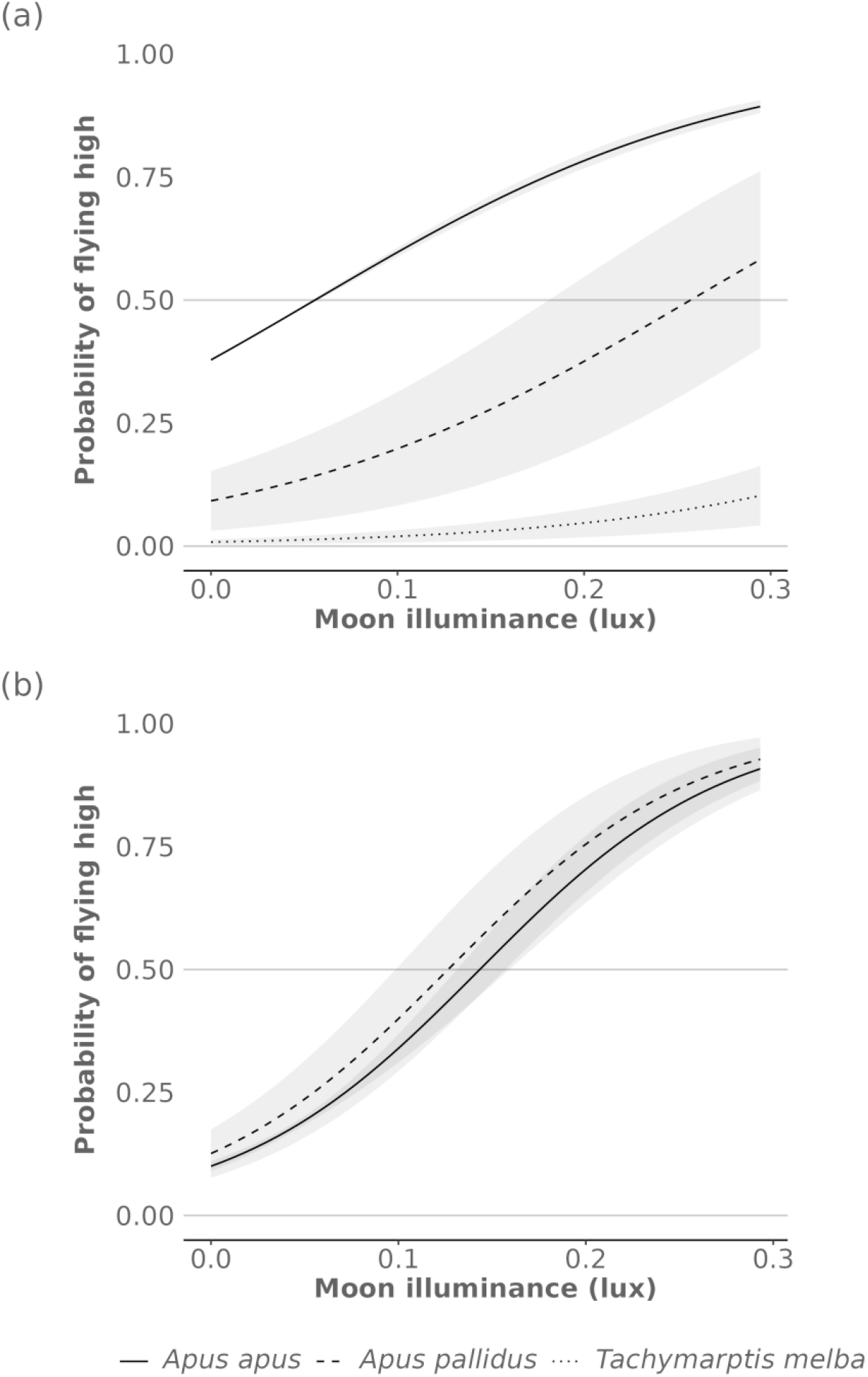
Marginal effect sizes by species for the generalised linear mixed effects logistic models relating moon luminance with flight height for both MDL, i.e. showing the probability of a high night-time flight relative to daytime flight altitudes for MDL **(a)** and GPS **(b)**. Grey 95% Confidence Intervals (CIs) were computed using a Wald z-distribution approximation and a standardised dataset.

**Table 1.**
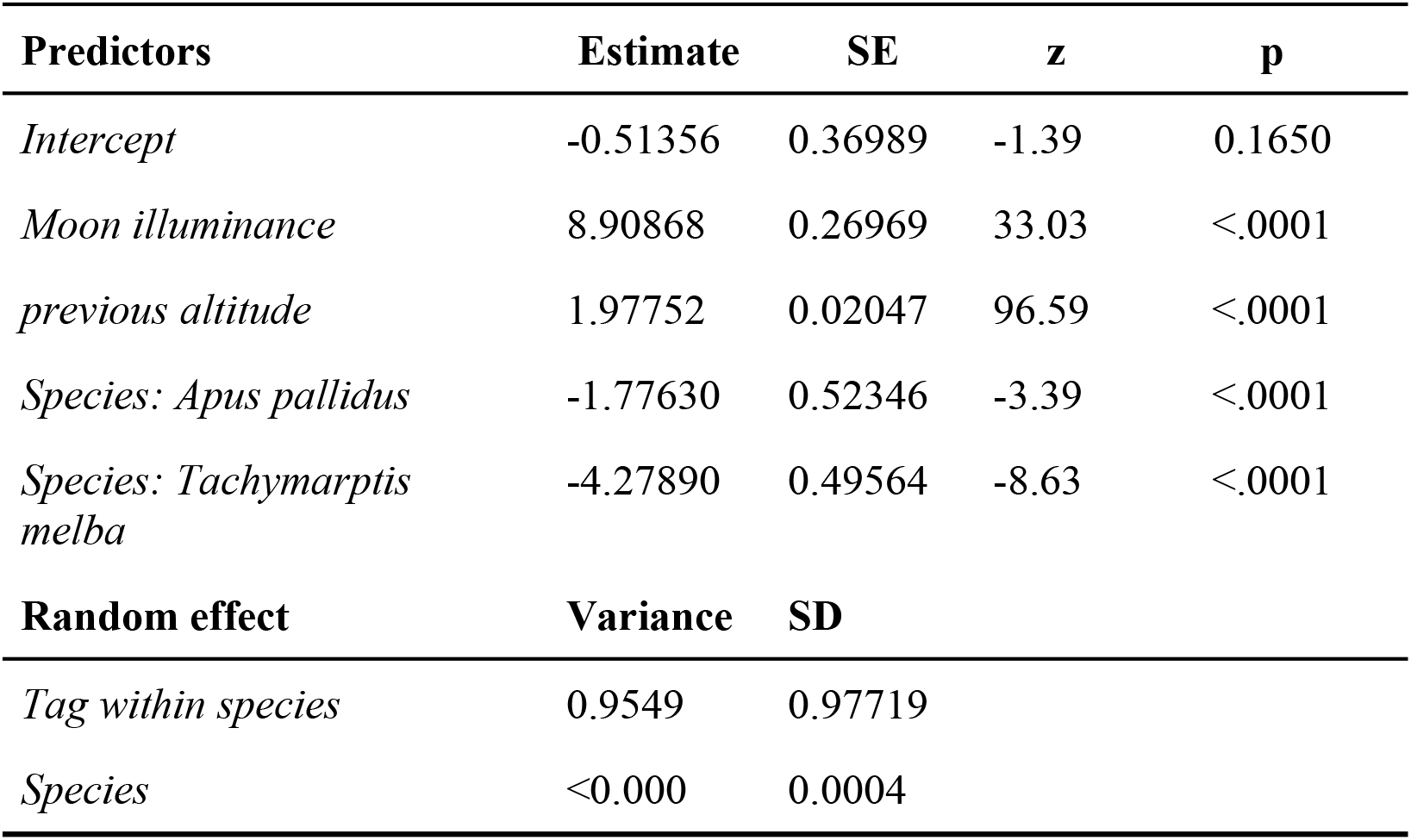
Model output of quantifying high or low flights in response to moon illuminance, using generalised linear mixed effects model and Multisensor Data logger (MDL) data. The model used moon illuminance, lagged altitude, to account for temporal autocorrelation, and species as fixed effects. The tag is nested within species as a random intercept to account for species and bird specific behaviour.

| Predictors | Estimate | SE | z | p |
| --- | --- | --- | --- | --- |
| *Intercept* | -0.51356 | 0.36989 | -1.39 | 0.1650 |
| *Moon illuminance* | 8.90868 | 0.26969 | 33.03 | <.0001 |
| *previous altitude* | 1.97752 | 0.02047 | 96.59 | <.0001 |
| *Species: Apus pallidus* | -1.77630 | 0.52346 | -3.39 | <.0001 |
| *Species: Tachymarptis melba* | -4.27890 | 0.49564 | -8.63 | <.0001 |
| Random effect | Variance | SD |  |  |
| *Tag within species* | 0.9549 | 0.97719 |  |  |
| *Species* | <0.000 | 0.0004 |  |  |

**Table 2.** Model output of quantifying high or low flights in response to moon illuminance, using generalised linear mixed effects model as measured from GPS data. The model used moon illuminance and species as fixed effects. The tag is nested within species as a random intercept to account for species and bird specific behaviour.

| Predictors | Estimate | SE | z | p |
| --- | --- | --- | --- | --- |
| *Intercept* | -2.1526 | 0.3991 | -5.392 | <.0001 |
| *Moon illuminance* | 15.3049 | 1.0566 | 14.485 | <.0001 |
| *Species: Apus pallidus* | 0.2165 | 0.4509 | 0.48 | 0.631 |
| Random effect | Variance | SD |  |  |
| *Tag within species* | 0.338 | 0.579 |  |  |
| *Species* | <0.000 | <0.000 |  |  |

Twilight ascends are especially pronounced for Common swifts with very brief (1h) long climbs up to an average maximum flight height of 2221 ± 629 m, and climbing on average 1807 ± 551 m (Figure 2d, Figure 3c). Twilight flights for Pallid and Alpine swifts are generally lower, only rising to average maximum flight heights of ~1500 m, and climbing ~1000 m (average maximum flight height: 1537 ± 656 m and 1464 ± 534 m, average flight height increase of 1187 ± 633 m and 897 ± 278 m, for Pallid and Alpine swifts respectively). When testing if nocturnal light affects a shift towards night-time away from civil twilight (for dusk and dawn) using a linear mixed effects model we find we found our model’s explanatory power weak (conditional R^2^ = 0.03, Table 3, Appendix Figure S9), with a statistical significant effect of both fixed effects, moon illuminance and twilight period (dusk/dawn).

**Table 3.** Model output of a linear mixed effects model comparing the timing of maximum twilight flight height (in minutes from civil twilight) in response to night-time moonlight illuminance using Multisensor Data Logger (MDL) data. Moon illuminance is a fixed effect with tags nested within species as a random intercept.

| **Predictors** | **Estimate** | **SE** | **t** | **p** |
| --- | --- | --- | --- | --- |
| *Intercept* | 8.562 | 1.141 | 7.507 | <.005 |
| *Moon illuminance* | 130.408 | 8.036 | 16.228 | <.0001 |
| **Random effect** | **Variance** | **SD** |  |  |
| *Tag within species* | 13.675 | 3.698 |  |  |
| *Species* | 1.977 | 1.406 |  |  |

For the Common and Pallid swifts, dusk marked the start of a period of sustained high activity continuing throughout the night until dawn with peaks during both twilight periods (Figure 3b). A formal test comparing daytime with night-time flight activity shows a marginal R^2^ of 0.7 for fixed effects alone. Where, for both the Common and Pallid swift the interaction effect between time-of-day (day vs. night) and species is statistically significant and positive, this interaction is statistically significant and negative for Alpine swifts (Appendix Table S6, Figure S9).

## Discussion

Several aerial insectivores have shown to profit from night-time illuminance conditions, either from artificial sources or through moonlight. These favourable illumination conditions allow for prolonged foraging behaviour. Direct observations of Common swifts show documented prolonged flight activity in the presence of intense urban light pollution (Amichai & Kronfeld-Schor 2019), while it has been hypothesised that Northern Black swifts (*Cypseloides niger borealis*) use favourable night-time light conditions to continue foraging (Hedenström *et al*. 2022). In this study we explored if three western palearctic swift species, the Common, Pallid and Alpine swift, show divergent night-time behaviour across their African non-breeding season in response to moonlight illuminance.

Our results show that moonlight driven night-time responses are strongly species and context dependent. We note the strongest response (absolute flight heights and moonlight illuminance sensitivity) in the Common swift, a reduced response in Pallid swifts and no response in Alpine swifts. Species responses fall along a gradient in body and prey size, and previously described flight patterns (Collins *et al*. 2009, Helms *et al*. 2016). We show a weak relationship between night-time illuminance driven responses and twilight ascending behaviour across all species, suggesting a decoupling of both behaviours, separating night-time foraging from social, zeitgeber and orientation focussed behaviour.

### night-time moonlight response

Our study found three distinct night-time flight patterns in response to changing moonlight illuminance. This varying sensitivity in the probability of a high night-time flight suggests that the magnitude and frequency of night-time moonlight responses are highly species and context specific.

In Common swifts, the strong synchrony of nocturnal flight behaviour with moon illuminance showed a pronounced non-linear relationship in their flight height response, with a low threshold value relative to their mean daytime flight behaviour (Figure 2d, Figure 4a). This behaviour was limited in flight height (Figure 2d, Figure 3d), but still allowed the Common swifts to rise more than 1500 m. Little change in moonlight is therefore required to trigger a change in their flight height. Research on the American Black swift has shown that responses to decreasing moon illuminance, in response to a moon eclipse, are indeed fast and reversible (Hedenström *et al*. 2022). Our analysis corroborates these observations where moonlight synchronised flight patterns are at times abruptly interrupted, only to be resumed later in the night (Figure 3a, months of January and February). Conversely, during the new moon, in absence of any moonlight, flights remained consistently low. For Pallid swifts, higher illuminance values are required to trigger a higher flight (Figure 4a-b) as shown by our model results of MDL data.

Less pronounced moonlight driven behaviour in Pallid swifts is likely due to common low altitude coastal foraging behaviour around sea breeze fronts (Kearsley et al. 2022) which limits the ability to rise to greater heights. Here, offshore winds at higher altitudes (>1000m) would push birds away from land. GPS data shows that Common swifts rarely position themselves near or over coastal waters, as shown by only 14 registered locations, while almost a quarter of the positions of the Pallid swifts are coastal foraging locations (Kearsley et al. 2022, Appendix Table S2). This suggests that moonlight illuminance driven responses are mediated by synoptic and macroscale weather. In contrast, inland continental positions showed higher night-time flights of Pallid swifts (Table 1 in Kearsley *et al*. 2022). It was speculated that these flights could be linked to common aggregation of insects at inversion layers at the dry side of the ITCZ (Reynolds *et al*. 2008, Nicholson 2009). Our current analysis puts these results in a different perspective, and shows a strong connection to moonlight illuminance. However, we can not exclude the influence of weather and seasonal changes in frequency of the moonlight illuminance responses for both the Common and Pallid swifts. Overview plots show that the response is weakest from December through February (Figure 3, Appendix S1 - S2). These time windows correspond with time spent in tropical regions where cloud conditions or large frontal systems might obscure or influence moonlight driven flight responses. We suggest that clouds and a swift’s geographic position with respect to seasonal (local) weather, might limit moon illuminance responses in more cloudy tropical areas.

In contrast to the behaviour of both the Common and Pallid swift, the Alpine swift does not show a moonlight illuminance driven response in flight height and activity (Figure 3a - b, Figure 4a). Despite being a small aerial insectivore, which is continuously on the wing (Meier *et al*. 2018), they are twice the size of both the Common and Pallid swift, weighing ~100g compared to ~40g (Collins *et al*. 2009). Body size positively correlates with prey size across seven species of swifts, with the Alpine swift targeting consistently larger prey than *Apus sp.* (Collins *et al*. 2009). It is a well observed phenomena that insects increase their flight height and activity during moonlit nights (Brown & Taylor 1971, Bowden & Gibbs 1973, Danthanarayana 1986) and insect size distributions are strongly vertically stratified based, with small insects flying higher than large ones (Helms *et al*. 2016, Jansson *et al*. 2021). As such, Alpine swift, preferring larger (low flying) insects, would naturally forage at lower altitudes than the Common and Pallid swift which prefer high flying smaller insects. We show that cyclical moonlight driven increases in flight height in search for prey seems plausible, where swift flight patterns follow a size dependent continuum in (insect) prey size (Helms *et al*. 2016, Jansson *et al*. 2021).

While moonlight determines flight height in Common and Pallid swifts, it does not affect flight activity. During all nights flight activity remained high, suggesting more intense flapping activity and conversely less soaring flights (Figure 3b). In contrast, Alpine swifts do not show sustained high activity during the night, with activity values similar to those during the day (Figure 3b). The lack of thermals during night-time might explain some of this dynamic, although the contrasting behaviour of the Alpine swift puts this hypothesis in question. A strong moonlight driven decrease in activity as mentioned by Hedenström *et al*. (2022) is missing in all our observations (Appendix Figure S3-S5).

Although the spatial distribution of swift species seem to follow a size dependent vertical distribution of insects during the night we can not exclude predator avoidance as part of the moonlight illuminance driven response. The Eleonora’s falcon shares overlapping regions with all three species during the non-breeding species (Vansteelant *et al*. 2021), and is known to hunt at night under favourable artificial or moonlight conditions (Buij & Gschweng 2017). Escaping from falcons by rising flights is a common predator evasion strategy in swifts (Hedenstrom 2001). Giant noctule bats (*Nyctalus lasiopterus*) have been shown to feed upon passerine birds during migration (Popa-Lisseanu *et al*. 2007, Ibáñez *et al*. 2016). Noctule bats show considerable flexibility in foraging timing and space use in response to food availability. However, contrasting flight strategies and/or prolonged periods of high flights across swifts would make this scenario less likely.

Furthermore, the observed patterns question the notion that birds roost throughout the night, on the wing (Rattenborg *et al*. 2016). Nocturnal moonlight driven changes in flight height and activity for the Common and Pallid swifts suggest that, at least, these species adapt their waking flights to the light environment. This suggests plasticity in their sleep-wake cycle, where it has been suggested that descending twilight flights might serve as a time for rest (Hedenström 2016, Hedenström *et al*. 2022).

### Twilight behaviour

For all species, we find pronounced twilight ascents aligning with a period of high flight activity. For both the Common and Pallid swifts this high activity continues through the night regardless of light driven changes in flight height (Figure 2a-b). In contrast, Alpine swifts see isolated peaks during twilight, with activity values similar to daytime values during the night.

Maximum twilight flight heights are attained before civil dusk or right after civil dawn (Figure 4). Given our current data, we do not show strong support for a shift in twilight flight timing in relation to moon illuminance. While it would be energetically favourable to not descend when high moonlight illuminance conditions follow later in the night (or conversely moonlight illuminance is followed by twilight conditions) we do not observe such behaviour. Swifts do not seem to anticipate a future moonlit night or coming twilight conditions (Figure 2b, Figure 3a). All swifts descend after their twilight ascent, even briefly, before any moonlight illuminance response. Due to the short duration of twilight events, the sampling rate and the uncertainty on twilight events linked to positional accuracy the statistical power of our analysis is limited.

The full scope of the twilight behaviour, and its connection to night-time activity, remains poorly understood. It has been shown that Pallid swifts overtop sea-breeze fronts using higher twilight and night-time flights in order to retain a favourable foraging position (Kearsley *et al*. 2022). Therefore, an alternative hypothesis exists for at least a part of the non-breeding season of Pallid swifts, while for other species other factors can not be excluded. A weak moonlight illuminance sensitivity, moving into and out of twilight conditions does not exclude the re-orientation hypothesis (Muheim 2011, Muheim *et al*. 2018) with flights linked to the timing of maximum sunlight polarisation. The most pronounced twilight ascends are recorded for the Common swift. This species crosses the equator and would benefit the most from a twilight recalibrated compass (Cochran *et al*. 2004). Strong twilight ascending flights could suggest re-orientation events. However, past analysis of Alpine swift data showed social bonding through screaming parties dominated over re-orientation (Meier *et al*. 2018). Foraging behaviour can not be fully excluded either. Both moonlight illuminance (Brown & Taylor 1971, Bowden & Gibbs 1973, Danthanarayana 1986) and twilight conditions have been linked to increased insect activity of small swarming insects such as mosquitoes, flies and ants (Jansson *et al*. 2021), making up a sizeable proportion of the diet of small aerial insectivore (tropical) bats (Racey & Swift 1985, Pavey *et al*. 2001). During the breeding season Common swifts have been observed to prey on swarming insects at twilight near the Afsluitdijk, The Netherlands (Dokter *et al*. 2013). However, limited observational evidence exists of such behaviour during the non-breeding season. Finally, due to the continued active behaviour in moonlit nights the hypothesis of descending twilight flights as a resting phase (Hedenström 2016, Hedenström *et al*. 2022) needs consideration as this behaviour is pronounced and consistent across all three permanently airborn swift species (Liechti *et al*. 2013, Hedenström 2016, Meier *et al*. 2018, Hedenström *et al*. 2019).

In the assessment of both the night-time moonlight illuminance responses and twilight behaviour there is a need for co-located measurements of insect prey to confirm a distinct foraging aspect over potential social, sleep, orientation, or predator avoidance based behaviour. The co-occurence of swift along an insect prey size dependent distribution, following behaviour of insects as described in literature, would call into question if, like nightjars (Jetz *et al*. 2003), at least the Common and Pallid swifts should be considered crepuscular-nocturnal aerial insectivores.

## Conclusion

We show that both the Common and Pallid swifts engage in night-time ascending behaviour in the presence of moonlight, while no such behaviour is observed in Alpine swifts. These flight patterns follow a weight to prey-size vertical stratification of insects as described in the literature. Our analysis changes the interpretation of the diurnal movements of night-time ascents of Pallid swifts previously described, and suggests similarities between both Common and Pallid swift behaviour and the behaviour observed in both the Northern Black swift and nightjars. All these species seem to extend their night-time foraging behaviour in the presence of moonlight. As all swift species considered (so far) have documented continuous flights during the non-breeding season, we suggest swifts optimise their flight behaviour to adapt to favourable night-time light conditions, weather conditions and a size dependent (vertical) insect prey distribution.

## Acknowledgements

Belora vzw, Locus Developments bv (Gent) and the Municipal government of Vila Nova de Famalicão funded GPS loggers. We are grateful for the support by Migrate Technology Ltd. for providing prototype Multisensor Data Loggers in a research and development collaboration. We thank PathTrack Ltd for their custom data processing, and discussions regarding data quality. Moon phase icons were taken from the Emoji One BW v2.0 library from JoyPixels (https://joypixels.com/).

## Ethics

Swiss loggers were attached under a ringing licence of the Federal Office for the Environment FOEN and ethical approval of the veterinary office of canton of Aargau under the licence 26552 / LU0415. All research at the Arrabida/Famalicão sites was carried under all the required legal requirements of the ICNF (Portuguese Institute for the Conservation of Nature and Forests), ringing licences 134/2021 and 144/2022. For research at the Gent Voorhaven site the licence was issued by the Agency for Nature and Forest, Belgium (Flanders) number ANB/BL-FF/VERG/11-00316.

## Code & Data availability

The manuscript’s database and code supporting our findings is made available on Github <bluegreen-labs.github.io/swift_lunar_synchrony/> and a matching Zenodo Digital Repository <https://doi.org/10.5281/zenodo.7814214> under a CC-BY 4.0 licence.

