## Appendix for "Moonlight synchronous flights across three western palearctic swifts mirror size dependent prey preferences"

#### Methods

Swift movements were categorised as migration (corridor movements) and stationary (area-restricted search) using a two-state Hidden Markov Model (McClintock & Michelot 2018, HMM), with missing positions filled using continuous-time correlated random walk data (Johnson *et al.* 2008) to create a homogeneous six-hour time series. Step distances (i.e. Euclidean distances between subsequent GPS positions) were modelled using a gamma distribution, while turning angle distributions followed a von Mises distribution (initial parameters; gamma distributions: μf = 50 km, σf = 10 km; μm = 80 km, σm = 10 km; von Mises distributions: μf = 0, κf = 1; μm = 0, κm = 2, migration and foraging denoted with subscripts m and f, respectively). State classes (i.e. migration, stationary) were returned using global encoding with the Viterbi algorithm (Zucchini *et al.* 2016, McClintock & Michelot 2018).

MDL atmospheric pressure recorded was converted to altitude (z) using the international standard atmosphere which is defined as:

$$z = \frac{T_0}{L}\left(\left(\frac{P_0}{P}\right)^{\frac{LR_0}{g}} - 1\right)$$

With $T_0$ the temperature at sea level (288.15 K), L the temperature lapse rate (-0.0065° K $m^{-1}$), $P_0$ the standard atmospheric pressure at sea level (1013.25 hPa), P the measured air pressure (hPa), g the gravitational acceleration (9.81 m $s^{-1}$) and $R_0$ the universal gas constant (287.053 J $kg^{-1}$ $K^{-1}$). The Migrate Technology Intigeo-CAR and Swiss Ornithological

Institute GDL accelerometer read bursts of 32 x-axis (pitch, zero is horizontal) and 32 z-axis samples at 50 Hz. The sampling bursts are set so that wing beat frequency from 1.6 Hz to 25 Hz should be validly recorded to avoid aliasing. Activity values reported are the sum of consecutive differences recorded (up down movements) or peak g-forces applied over the set measurement time interval (either 30 or 5 min, for Migrate Technology Intigeo-CAR or Swiss Ornithological Institute GDL sensors respectively).

Additional summary statistics to those described in the main manuscript are provided. We report mean average flight altitude and height, activity with their uncertainty (on standard deviation). Furthermore, we tabulate current and/or long term swift recapture rates.

### Tables

**Table S1.** Overview table of the Multisensor Data Logger (MDL) meta-data. We report the logger tag number, the sun angle used in geolocation calculations, the start and end date defining the maximum available data for the non-breeding foraging season, and the variables logged. Variables logged are denoted with T, Pr, Act, P, Alt for temperature, Pressure, Activity, Pitch and Altitude, respectively. All Multi Data Loggers (MDLs) register light levels, while all GPS units register altitude in addition to position.

| Species | Site | Tag | Season | Sun angle | Start date | End date | variables |
|---|---|---|---|---|---|---|---|
| *Apus apus* | Gent Voorhaven | CC874 | 2021 | -6 | 2021-08-16 | 2021-12-19 | T, Pr |
| | | CC875 | 2021 | -5 | 2021-08-15 | 2022-05-06 | T, Pr |
| | | CC876 | 2021 | -5 | 2021-08-15 | 2022-04-29 | T, Pr |
| | | CC877 | 2021 | -7 | 2021-08-15 | 2022-04-29 | T, Pr |
| | | CC893 | 2021 | -7 | 2021-08-15 | 2022-05-05 | T, Pr, Act, P |
| | | CC895 | 2021 | -5 | 2021-10-01 | 2022-04-28 | T, Pr, Act, P |
| | | CC896 | 2021 | -7 | 2021-09-25 | 2022-04-28 | T, Pr, Act, P |
| *Apus pallidus* | Arrabida | CC883 | 2021 | -5.5 | 2021-09-18 | 2022-03-09 | T, Pr |
| | | CC902 | 2021 | -7.5 | 2021-09-15 | 2022-03-11 | T, Pr, Act, P |
| | Famalicão | CC904 | 2021 | -7 | 2021-09-27 | 2022-05-16 | T, Pr, Act, P |
| | | CC886 | 2021 | -6.5 | 2021-11-01 | 2022-04-09 | T, Pr |
| | | CC906 | 2021 | -4.5 | 2021-10-10 | 2022-04-14 | T, Pr, Act, P |
| | | CC907 | 2021 | -6.5 | 2021-10-15 | 2022-04-09 | T, Pr, Act, P |
| | | CC899 | 2021 | -7.5 | 2021-11-03 | 2022-03-23 | T, Pr, Act, P |
| *Tachymarptis melba* | Baden | 24SW | 2019 | -6.5 | 2019-10-01 | 2020-03-15 | T, Pr, Act, P |
| | | 24MP | 2019 | -6.5 | 2019-10-01 | 2020-03-15 | T, Pr, Act, P |
| | | 24SK | 2019 | -7 | 2019-10-01 | 2020-03-15 | T, Pr, Act, P |
| | | 24SI | 2019 | -7 | 2019-10-01 | 2020-03-15 | T, Pr, Act, P |
| | Pirasali | 22MA | 2018 | -7 | 2018-10-01 | 2019-03-15 | T, Pr, Act, P |
| | | 22LK | 2018 | -7.5 | 2018-10-01 | 2019-03-15 | T, Pr, Act, P |
| | | 22LE | 2018 | -7 | 2018-10-01 | 2019-03-15 | T, Pr, Act, P |
| | | 22LA | 2018 | -7 | 2018-10-01 | 2019-03-15 | T, Pr, Act, P |
| | | 22ME | 2018 | -6.5 | 2018-10-01 | 2019-03-15 | T, Pr, Act, P |

**Table S2.** Overview table of GPS meta-data. We report the logger tag number, the start and end date defining the maximum available data for the non-breeding foraging season, and the variables logged. All GPS units register altitude in addition to position.

| Species | Site | Tag | Season | Start date | End date | variables |
|---|---|---|---|---|---|---|
| *Apus apus* | Gent Voorhaven | Tag21900 | 2020 | 2020-08-26 | 2020-12-06 | Alt |
| | | Tag57085 | 2021 | 2021-08-20 | 2022-05-05 | Alt |
| *Apus pallidus* | Arrabida | Tag21417a | 2019 | 2019-08-26 | 2020-03-24 | Alt |
| | | Tag21453 | 2019 | 2019-08-28 | 2020-04-14 | Alt |
| | | Tag21482 | 2019 | 2019-08-23 | 2020-05-07 | Alt |
| | | Tag21472 | 2019 | 2019-08-27 | 2020-04-28 | Alt |
| | | Tag21460 | 2019 | 2019-08-27 | 2020-04-11 | Alt |
| | | Tag21417b | 2020 | 2020-09-12 | 2021-04-03 | Alt |
| *Apus pallidus* | Famalicão | Tag56597 | 2021 | 2021-09-19 | 2022-03-03 | Alt |
| | | Tag57069 | 2021 | 2021-09-23 | 2022-04-04 | Alt |

**Table S3.** Overview table of the GPS meta-data. We report the deployment site, logger tag number, the moon phase and the mean night-time flight height. All GPS units register position and altitude.

| Species | Site | Moon phase | Continental flight height (mean ± sd) | Coastal flight height (mean ± sd) | Surface elevation (mean ± sd) | N (continental / coastal) |
|---|---|---|---|---|---|---|
| *Apus apus* | Gent Voorhaven | New | 276 ± 244 | / | 521 ± 250 | 97 / 0 |
| | | First Quarter | 370 ± 404 | 350 ± 66 | 527 ± 330 | 110 / 2 |
| | | Full | 921 ± 684 | 270 ± 267 | 514 ± 381 | 113 / 12 |
| | | Last Quarter | 518 ± 555 | / | 529 ± 352 | 107 / 0 |
| *Apus pallidus* | Arrabida | New | 564 ± 481 | 452 ± 491 | 277 ± 136 | 281 / 121 |
| | | First Quarter | 713 ± 556 | 549 ± 574 | 274 ± 151 | 264 / 151 |
| | | Full | 1075 ± 516 | 841 ± 739 | 267 ± 141 | 278 / 126 |
| | | Last Quarter | 701 ± 492 | 530 ± 535 | 278 ± 138 | 246 / 120 |
| *Apus pallidus* | Famalicão | New | 444 ± 288 | 767 ± 521 | 263 ± 183 | 83 / 2 |
| | | First Quarter | 591 ± 285 | 592 ± 479 | 260 ± 194 | 61 / 13 |
| | | Full | 1075 ± 391 | 1007 ± 1240 | 277 ± 194 | 70 / 14 |
| | | Last Quarter | 617 ± 482 | 903 ± 749 | 270 ± 190 | 84 / 3 |

**Table S4.** Overview table of the Multisensor Data Logger (MDL) meta-data. We report the deployment site, logger tag number, the moon phase and the mean night-time flight altitude, flight height and activity level.

| Species | Site | Moon phase | Altitude (mean ± sd) | Activity (mean ± sd) | Flight Height (mean ± sd) | N |
|---|---|---|---|---|---|---|
| *Apus apus* | Gent Voorhaven | New | 978 ± 434 | 974 ± 656 | 332 ± 327 | 6798 |
| | | First Quarter | 1201 ± 733 | 953 ± 669 | 615 ± 699 | 7244 |
| | | Full | 1810 ± 954 | 1097 ± 643 | 1226 ± 909 | 7689 |
| | | Last Quarter | 1231 ± 714 | 1001 ± 647 | 649 ± 676 | 7374 |
| *Apus pallidus* | Arrabida | New | 746 ± 397 | 972 ± 663 | 401 ± 332 | 1630 |
| | | First Quarter | 804 ± 492 | 910 ± 675 | 421 ± 424 | 1683 |
| | | Full | 948 ± 563 | 1040 ± 671 | 606 ± 462 | 1878 |
| | | Last Quarter | 919 ± 466 | 1086 ± 636 | 523 ± 407 | 1762 |
| | Famalicão | New | 640 ± 360 | 800 ± 652 | 298 ± 311 | 4190 |
| | | First Quarter | 761 ± 481 | 819 ± 658 | 393 ± 421 | 4657 |
| | | Full | 974 ± 593 | 938 ± 650 | 634 ± 502 | 4562 |
| | | Last Quarter | 791 ± 477 | 906 ± 656 | 432 ± 421 | 4038 |
| *Tachymarptis melba* | Baden | New | 861 ± 282 | 30 ± 37 | 196 ± 183 | 3843 |
| | | First Quarter | 827 ± 294 | 28 ± 39 | 187 ± 168 | 4144 |
| | | Full | 928 ± 315 | 38 ± 48 | 253 ± 209 | 3828 |
| | | Last Quarter | 903 ± 299 | 26 ± 37 | 219 ± 201 | 3542 |
| | Pirasali | New | 1303 ± 366 | 31 ± 40 | 331 ± 264 | 3739 |
| | | First Quarter | 1261 ± 375 | 33 ± 42 | 342 ± 303 | 3893 |
| | | Full | 1382 ± 402 | 42 ± 47 | 406 ± 334 | 3343 |
| | | Last Quarter | 1300 ± 371 | 42 ± 48 | 319 ± 281 | 3533 |

**Table S5.** Overview of GPS and MDL logger recapture rates as compared to seasonal or historical long term values by Costa & Elias (1998, *). Note that for Gent Voorhaven only two GPS tracks covered the fixed sampling scheme required for this study.

| Sensor | Site | # deployed | # recaptured | % recaptured | Control group recaptures (%, reference year) |
|---|---|---|---|---|---|
| GPS | Gent Voorhaven | 7 | 4 | 57 | 47 (2021) |
| | Arrabida | 6 | 5 | 83 | 65* |
| | Famalicão | 6 | 3 | 50 | 52 (2021) |
| MDL | Gent Voorhaven | 9 | 7 | 78 | 47 (2021) |
| | Arrabida | 5 | 2 | 40 | 65* |
| | Famalicão | 10 | 5 | 50 | 52 (2021) |
| | Baden | 8 | 4 | 50 | 86 (2019) |
| | Pirasali | 26 | 11 | 42 | 37 (2018) |

**Table S6.** Model output of quantifying activity in relation to species and time-of-day (TOD, night / day). The tag is nested within species as a random intercept to account for species and bird specific behaviour. Reporting fixed effects only.

| Fixed effects predictor | Estimate | SE | t-value | p-value |
|---|---|---|---|---|
| *Intercept* | 694 | 183 | 3.78 | 0.999 |
| *Species: Apus pallidus* | -140 | 259 | -0.542 | 0.999 |
| *Species: Tachymarptis melba* | -648 | 259 | -2.5 | 0.999 |
| *TOD* | 312 | 9.22 | 33.8 | <.0001 |
| *Species: Apus pallidus:TOD* | 26.2 | 12.2 | 2.15 | <.0001 |
| *Species: Tachymarptis melba:TOD* | -324 | 11.1 | -29.1 | <.0001 |

### Figures

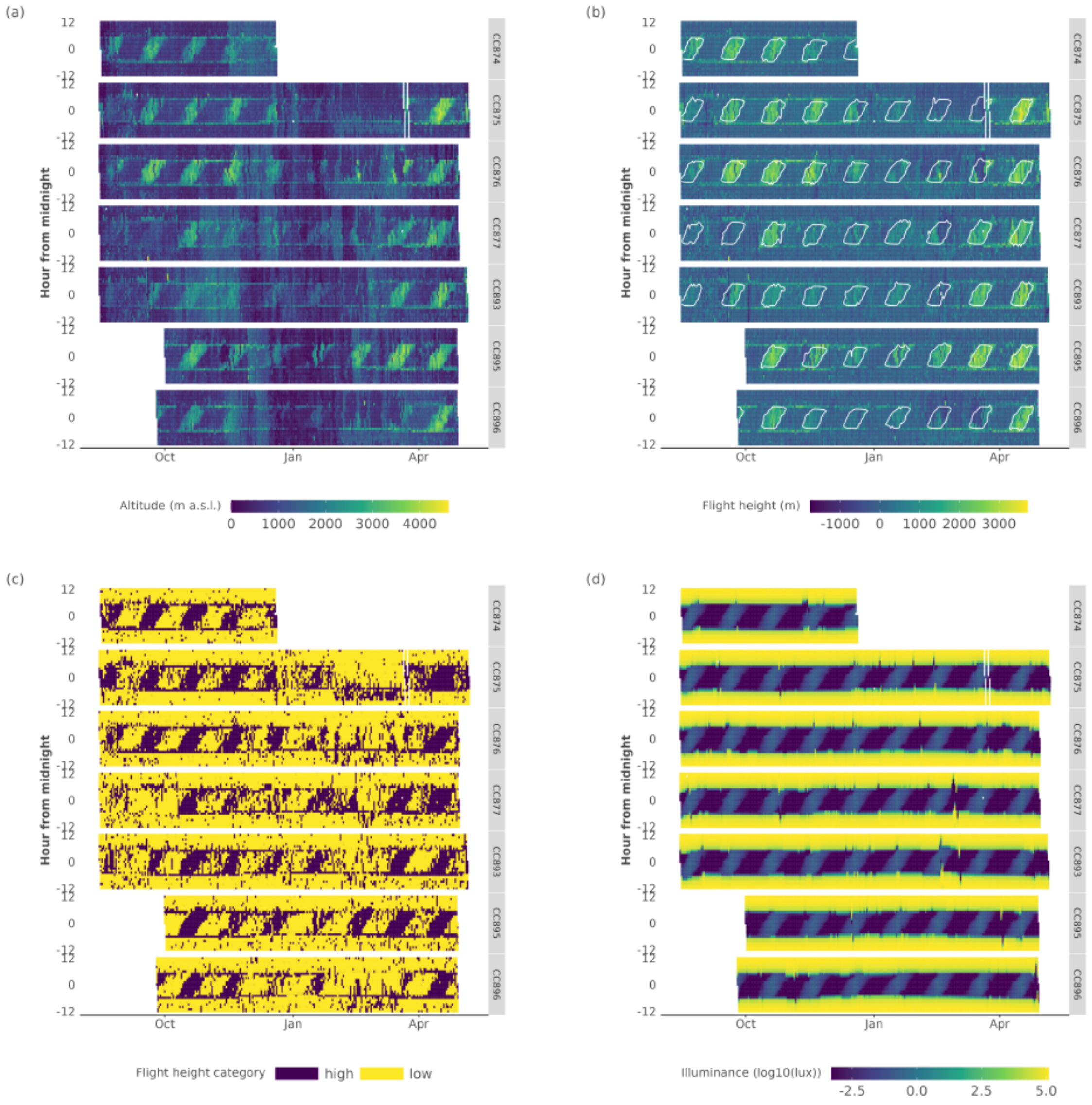


**Figure S1.** Overview plots (actograms) of *Apus apus* Multisensor Data Logger (MDL) data for flight altitudes (a), normalised flight heights (b) and the derived binary flight height classes (c). In addition, we provide the (log10) total illuminance for a given day (d). Data is shown for all MDL tags, with time centred on local midnight. Moon illuminance values outlining all values larger than 0.01 are contoured in panel (b).

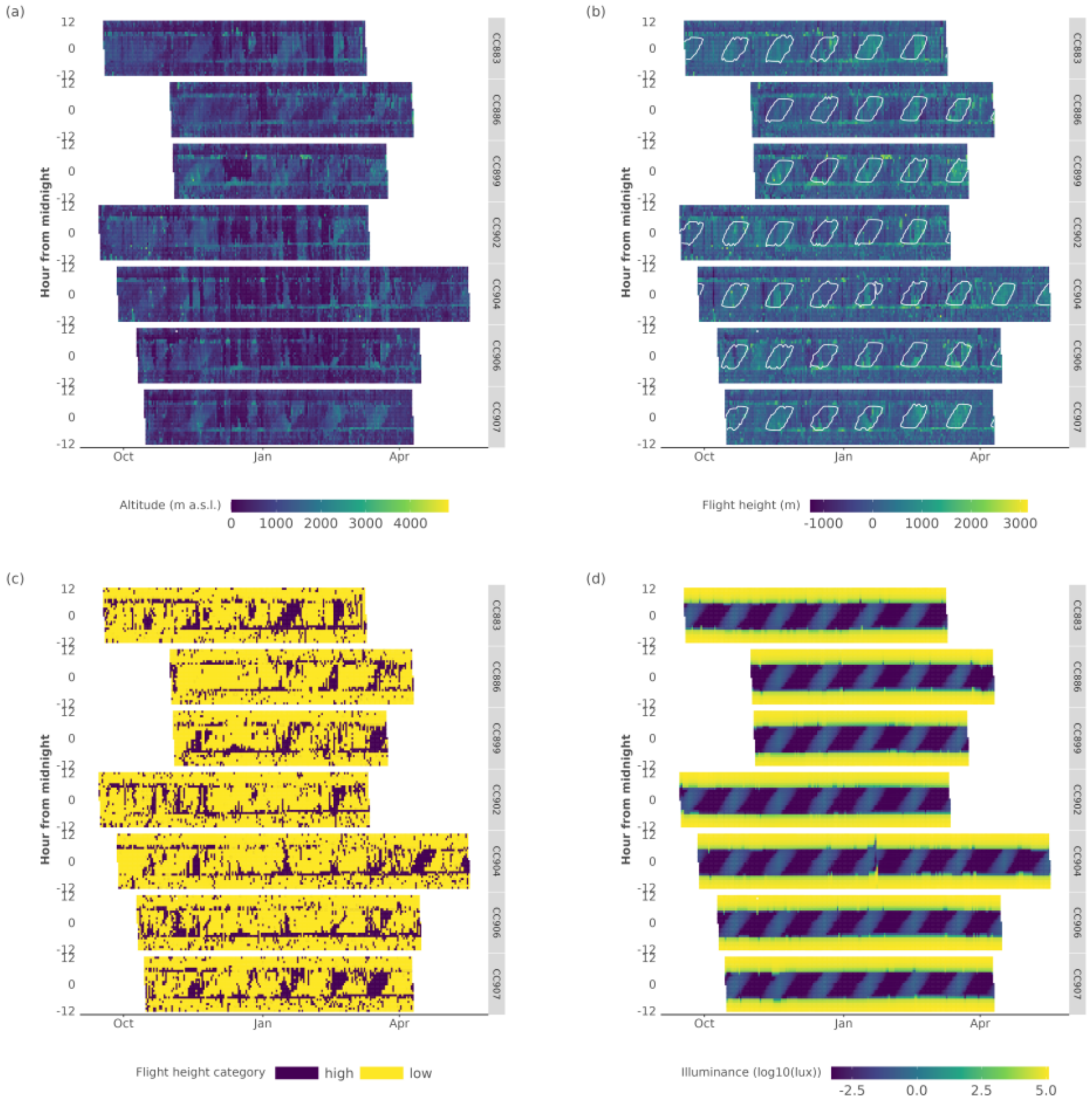


**Figure S2.** Overview plots (actograms) of *Apus pallidus* Multisensor Data Logger (MDL) data for flight altitudes **(a)**, normalised flight heights **(b)** and the derived binary flight height classes **(c)**. In addition, we provide the (log10) total illuminance for a given day **(d)**. Data is shown for all MDL tags, with time centred on local midnight. Moon illuminance values outlining all values larger than 0.01 are contoured in panel **(b)**.

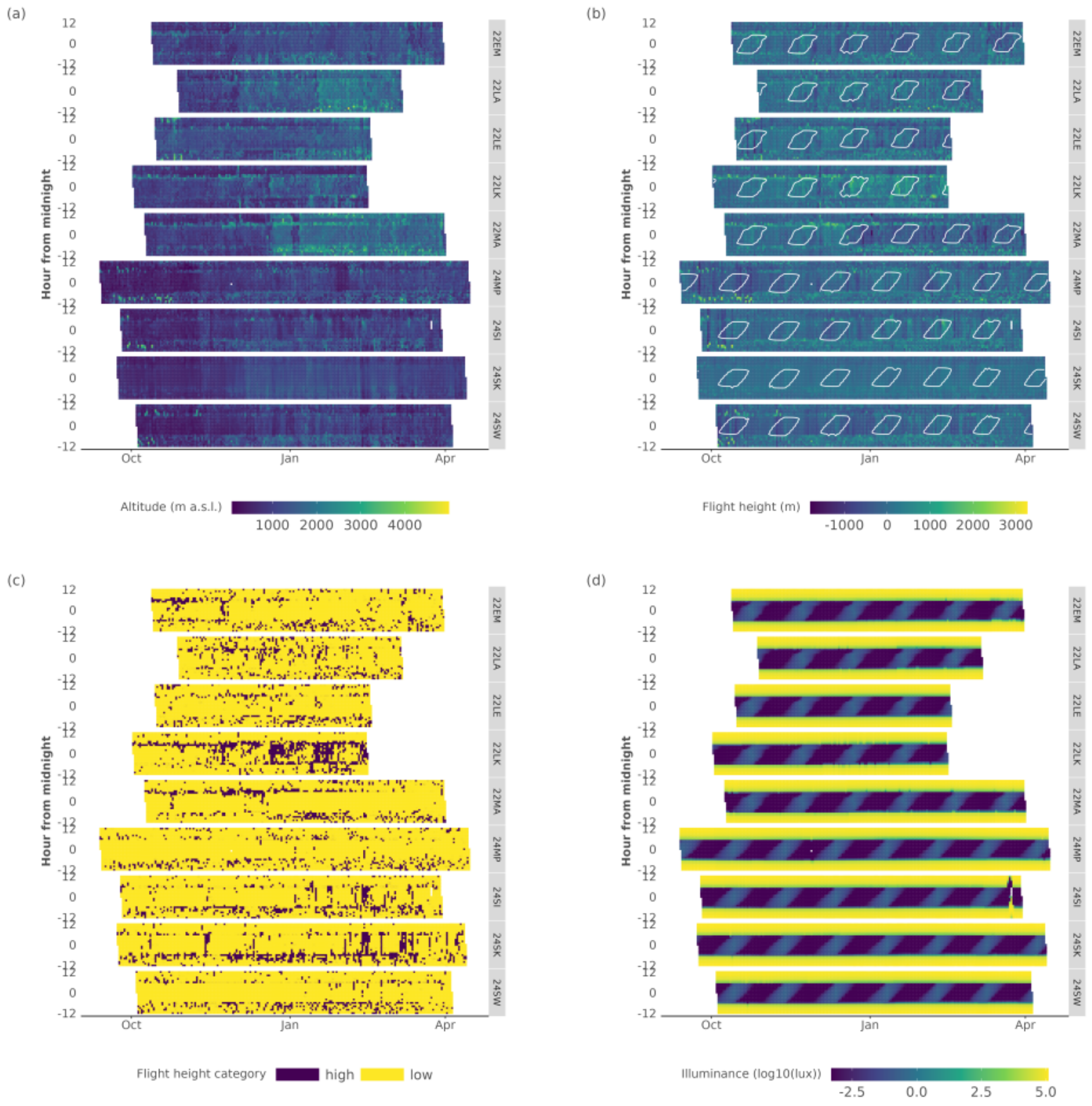


**Figure S3.** Overview plots (actograms) of *Tachymarptis melba* Multisensor Data Logger (MDL) data for flight altitudes **(a)**, normalised flight heights **(b)** and the derived binary flight height classes **(c)**. In addition, we provide the (log10) total illuminance for a given day **(d)**. Data is shown for all MDL tags, with time centred on local midnight. Moon illuminance values outlining all values larger than 0.01 are contoured in panel **(b)**.

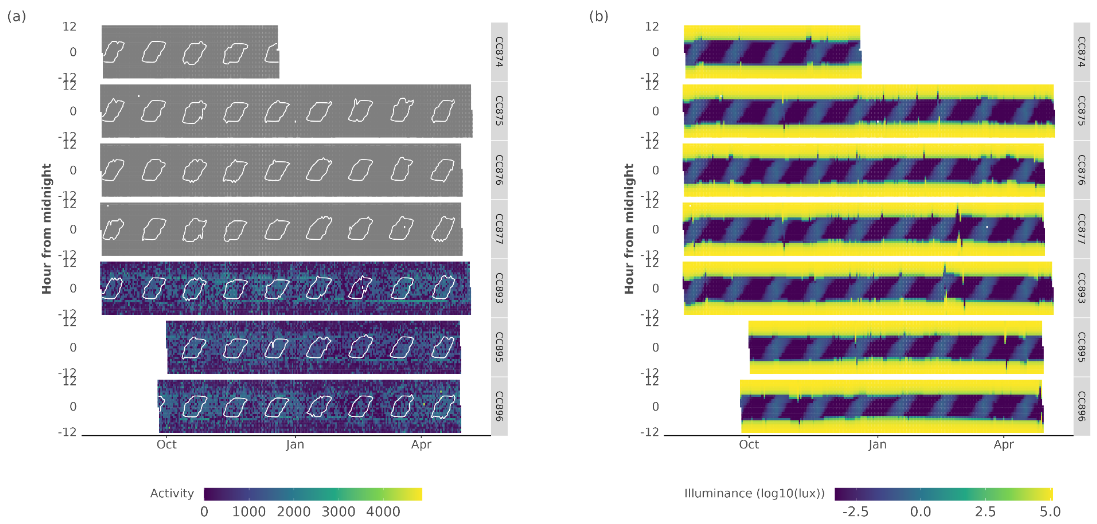


**Figure S4.** Overview plots (actograms) of *Apus apus* Multisensor Data Logger (MDL) data for flight activity (grey values are missing data) **(a)**. In addition, we provide the (log10) total illuminance for a given day **(b)**. Data is shown for all MDL tags, with time centred on local midnight. Moon illuminance values outlining all values larger than 0.01 are contoured in panel **(a)**. Missing values are shown as grey.

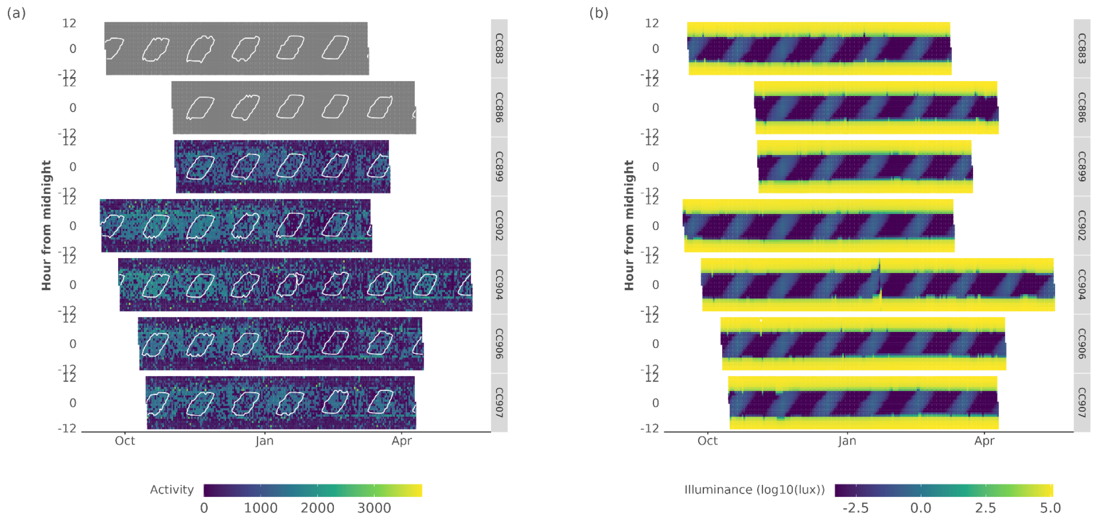


**Figure S5.** Overview plots (actograms) of *Apus pallidus* Multisensor Data Logger (MDL) data for flight activity (grey values are missing data) **(a)**. In addition, we provide the (log10) total illuminance for a given day **(b)**. Data is shown for all MDL tags, with time centred on local midnight. Moon illuminance values outlining all values larger than 0.01 are contoured in panel **(a)**. Missing values are shown as grey.

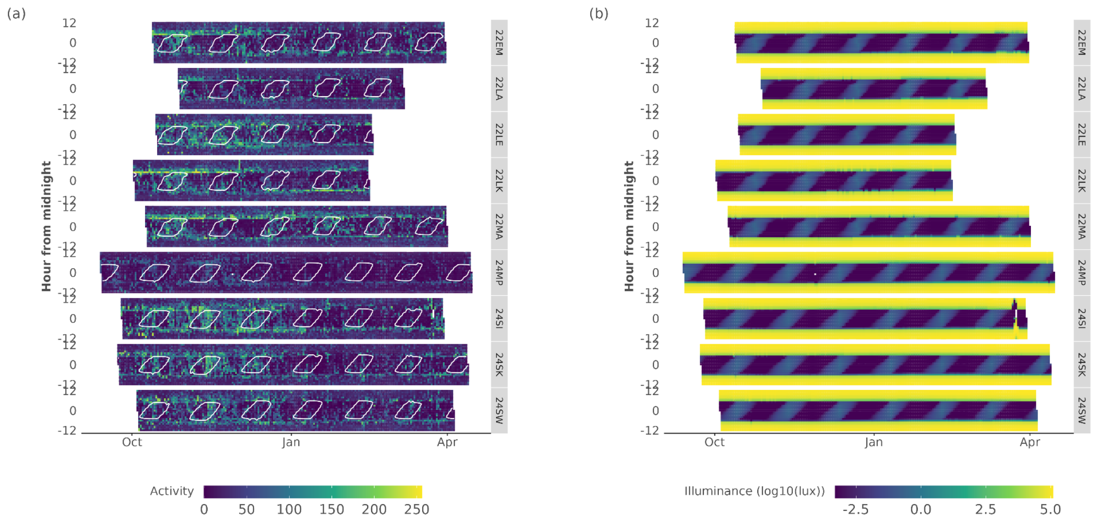


**Figure S6.** Overview plots (actograms) of *Tachymarptis melba* Multisensor Data Logger (MDL) data for flight activity (grey values are missing data) **(a)**. In addition, we provide the (log10) total illuminance for a given day **(b)**. Data is shown for all MDL tags, with time centred on local midnight. Moon illuminance values outlining all values larger than 0.01 are contoured in panel **(a)**. Missing values are shown as grey.

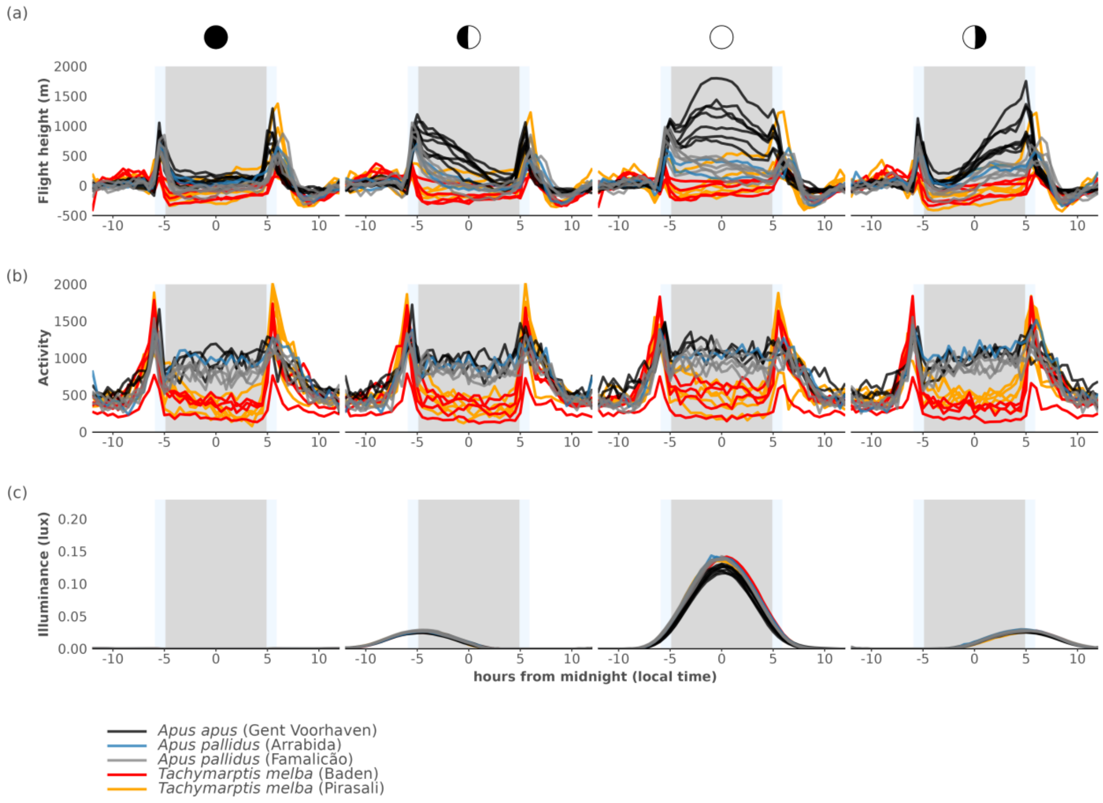


**Figure S7.** Average flight height (flight height in m, centred on a daytime mean of 0), the flight activity and derived moon illuminance (lux) centred on midnight during moon phases (a - c). Different sites and species are indicated by the colour of the full lines, different lines represent different tags. The grey rectangle in plots **(a - c)** indicate the average astronomical night, where light blue rectangles approximate twilight as mean dusk and dawn ± 1 sd, respectively.

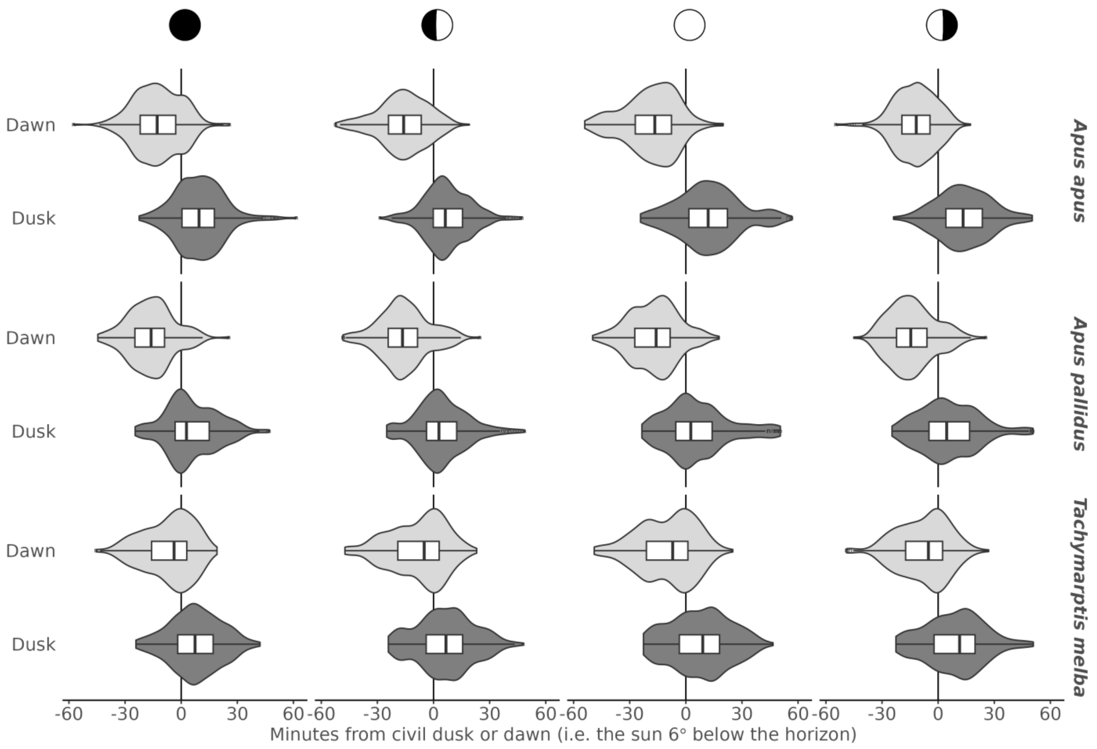


**Figure S8.** Box and violin plots of the distribution of the timing of maximum flight height during twilight, relative to civil dusk or dawn, by species and moon phase. The black vertical line at 0 indicates the timing of civil dusk or dawn. Values are expressed as minutes leading or lagging civil dusk or dawn events.

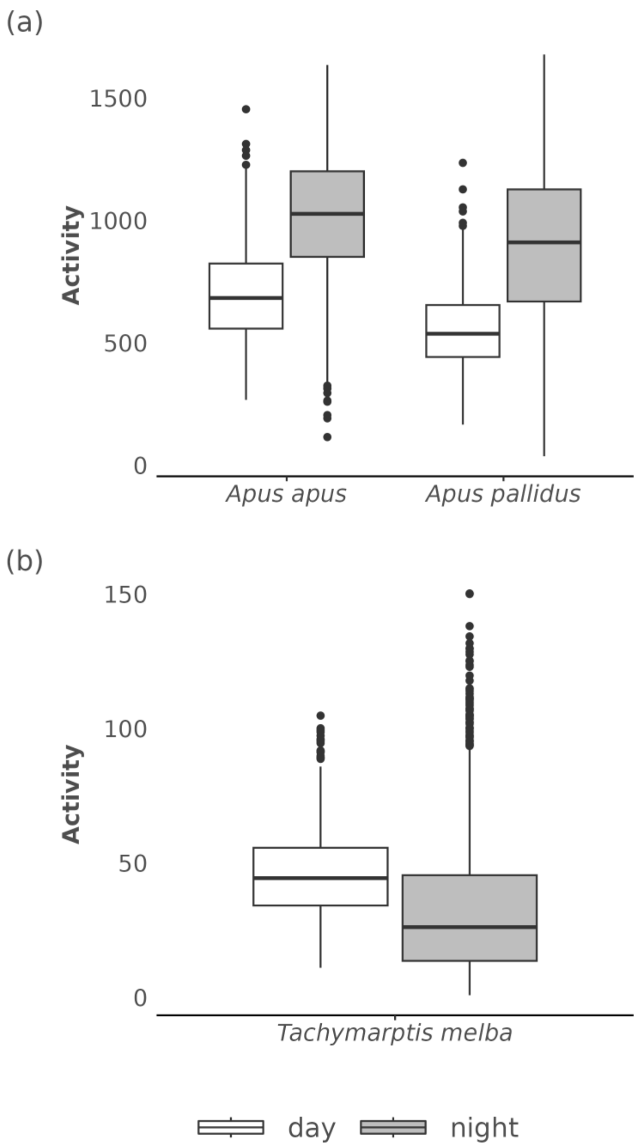


**Figure S9.** Box plots of the distribution of daytime and night-time flight activity by species.
